# A systemic approach allows to identify the pedoclimatic conditions most critical in the susceptibility of a grapevine cultivar to esca/Botryosphaeria dieback

**DOI:** 10.1101/2023.05.23.541976

**Authors:** Vinciane Monod, Vivian Zufferey, Matthieu Wilhelm, Olivier Viret, Katia Gindro, Daniel Croll, Valérie Hofstetter

## Abstract

Esca and/or Botryosphaeria dieback (esca-BD) are two of the most destructive grapevine trunk diseases in the world, disease complex which remains poorly understood. As some vine cultivars show highly variable susceptibility to esca-BD, we designed a four-year experiment to identify which environmental factors influence the expression of the disease. We collected epidemiological and physiological data once a year for four consecutive years in 19 vineyard plots located in four wine-growing regions of Western Switzerland. We compared these data with climatic data obtained from weather stations for these same plots for four years and over the long term. We also estimated the soil water holding capacity of each plot. Confounding factors were minimal because all vineyards were planted in 2003 with the same cultivar and all plants grafted in the same nursery with genetically homogeneous grafting material. Principal component and regression analyses of combined epidemiological, biotic and pedoclimatic data identified a positive correlation between soil water retention capacity and plant mortality due to esca-BD. These analyses also showed that leaf disease symptoms and apoplexy are more frequent when cold, wet periods are followed - or alternate with - hot, dry periods, and that apoplexy occurs more frequently when weather conditions change abruptly (cold, wet May followed by a hot June) and deviate significantly from long-term climatic conditions. Regression analyses show that the soil water holding capacity impacts less the disease expression when the climate is warm and dry, both at the regional and at year-specific levels. Having identified the most important environmental factors towards expression of esca-BD, this study allows recommendations to be given to the winegrowers for the cultivar studied but can also be used as a model to identify the environmental factors that influence the expression of fungal diseases in other grapevine cultivars, other grapevine trunk diseases and even in other woody plants.

## INTRODUCTION

Grapevine plants, with a lifespan of 30 years or more, are subject to a variety of biotic and abiotic stresses (Songy et al., 2019; Suzuki et al., 2014). Since the early 2000s, mature vineyards have begun to show increased yield and plant losses (Bertsch et al., 2013; Bertsch et al., 2009). This situation is attributed to the emergence of fungal diseases, linked to soil and climate factors (Bertsch et al., 2009; Bortolami, Farolfi, et al., 2021; Pastore et al., 2022; Songy et al., 2019). The cost of these diseases is high, involving numerous chemicals and/or biological fungicide treatments and replacement of dead plants (De Moura et al., 2012).

Among the emerging fungal diseases of grapevine, a complex called grapevine trunk disease (GTD) is primarily responsible for the death of young and mature grapevines worldwide (Claverie et al., 2020; Mondello et al., 2018). A very large number of new vineyards were planted during the 1990s, probably as a result of the spread of GTD in all countries where grapevines are grown (Gramaje & Armengol, 2011). The majority of these plants reached the age of disease expression at the same time, 7-10 years after their plantation, making the problem more noticeable (Gramaje et al., 2018). In addition, the generalisation of mechanical pruning has greatly increased the number of pruning wounds, the main entry point for the fungi associated with GTD (Makatini et al., 2014). These diseases pose a real threat to the viability and sustainability of viticulture (Bertsch et al., 2013; Kenfaoui et al., 2022).

The adult plants’ most destructive GTD worldwide are esca, Botryosphaeria dieback (Úrbez-Torres & Gubler, 2011), Eutypa dieback (Kuntzmann et al., 2010; Rolshausen et al., 2008; Trouillas & Gubler, 2010) and, to a lesser extent, Phomopsis dieback (Úrbez-Torres et al., 2013). All these diseases generate necroses in the grapevine trunk as well as foliar and/or shoot symptoms. Grapevine plants often suffer from more than a single GTD, due to their longevity and the multiple opportunities for fungal pathogens to infect wood mostly pruning (Bruez et al., 2014; Del Frari et al., 2019; Gramaje et al., 2018; Hofstetter et al., 2012). GTD infected plants may express foliar symptoms for several years and not always consecutively, but they usually die within a few years after the first disease symptoms are expressed (Bruez et al., 2013; Kenfaoui et al., 2022). All GTD-associated fungal species tested to date produce phytotoxic compounds (Andolfi et al., 2011; Martos et al., 2018; Masi et al., 2018). However, because these fungi live exclusively in wood, they have never been isolated from leaves (Bertsch et al., 2013; Bortolami et al., 2019), foliar symptoms are thought to result from the translocation of these phytotoxic compounds from the trunk to the leaves via sap flow (Bortolami, Gambetta, et al., 2021). To date, GTD foliar symptoms have rarely been reproduced under controlled conditions (Reis et al., 2016). The etiology of GTD pathogenic fungi remains poorly understood (Bertsch et al., 2013; Claverie et al., 2020; Mondello et al., 2018).

Soil and climate factors have also been suspected since long to influence GTD expression, particularly esca (Dubos et al., 2002). Environmental constraints seem to play a key role as they impact grapevine physiological behaviour by influencing host-microbiome interactions (Delmas, 2021), trigger the transition from a fungal endophyte state to a pathogenic state (Porras-Alfaro & Bayman, 2011; Saikkonen et al., 2003), and/or accelerate the translocation of metabolic compounds throughout the plant via sap flow (Claverie et al., 2020). Soils with high water holding capacity (deep, clayey soils) and climatic variations during the summer are factors reported to increase the risk of sap flow disruption (Surico et al., 2006). Vines with even mild foliar symptoms already suffer from hydraulic failure, a decrease in conductance in petioles and shoots, compared to healthy plants (Ouadi et al., 2019). Other important factors influencing disease expression appear to be linked to the plant physiology including the plant water status during growth periods, vigour (leaf area), carbon/nitrogen ratio in plant tissues (Berger et al., 2007) and functioning of the vascular system (Andreini et al., 2009; Edwards, Pascoe, et al., 2007). Two main hypotheses have been put forward to explain conductance failure: gas embolism (Canny, 1997) and/or occlusions (Sun et al., 2007), the latter either related or not with fungal pathogens. A recent study has shown that occlusions by tyloses and gels, probably induced remotely by phytotoxins produced by esca-associated fungi, lead to hydraulic failure in veins of esca-symptomatic leaves (Bortolami et al., 2019). However, the role of esca-BD associated fungi in the expression of foliar symptoms remains to be proven. Another study recently tested the effect of drought on grapevines under controlled conditions (Bortolami, Gambetta, et al., 2021) and found that drought prevents the expression of esca symptoms.

Numerous studies have attempted to understand the impact of soil type, particularly its ability to retain water, and/or climate on esca-BD disease expression (Bortolami, Farolfi, et al., 2021; Calzarano et al., 2018; Dubos et al., 2002; Fischer & Peighami-Ashnaei, 2019; Guérin-Dubrana et al., 2013; Marchi et al., 2006; Surico et al., 2000). However, such studies have been conducted in different regions, countries, or continents, often on different grape varieties with different disease susceptibility (Andreini et al., 2014), with plants of different ages, grown and pruned in different ways (Gramaje et al., 2018). The disparity of these studies has made it difficult to generalize the results obtained and to precisely identify the role of the pedoclimatic and biotic factors responsible for the variability in the incidence of esca-BD observed in different regions and for different grape varieties (Bertsch et al., 2009). Therefore, the impact of individual and/or combined pedoclimatic on the expression of esca-BD is not yet clearly established. While differences in esca-BD susceptibility between cultivars are relatively well documented (Bruez et al., 2013; Chacón-Vozmediano et al., 2021; Marchi, 2001; Serra et al., 2021), variation in esca-BD susceptibility of a single grape variety across multiple viticultural regions has rarely been tracked in long-term epidemiological studies (Dewasme et al., 2022; Guérin-Dubrana et al., 2013) and never, to our knowledge, by combining epidemiological data not only with climatic conditions but also with soil characteristics and plant physiological data.

To determine the impact of soil, climate and biotic factors on esca-BD expression, we reduced the effects of confounding factors by using a network of vineyards and studied for GTD expression since planting in 2003. In this network, 19 plots of an esca-BD susceptible cultivar (Gamaret) were selected, with all plants grafted with homogeneous scion and rootstock material in the same nursery and managed under uniform cultural practices. Such a network of vineyards can serve as a model to study the relationship between esca-BD expression and soil and climatic conditions. Since climatic demand and soil characteristics are varying within this network, this creates optimal conditions to study the influence of pedoclimatic factors on the prevalence of esca-BD. This experimental design allowed us i) to examine the variability of esca-BD incidence in vineyards located in different geographical areas, this independently of plant age, cultivar and vineyard cultural practices; ii) to explore the relationship (independence or correlation) between esca-BD expression, pedoclimatic factors and vine physiological factors in different vine-growing regions iii) to identify which factor(s) could be good candidate(s) to predict esca-BD expression.

## MATERIALS AND METHODS

### Study site, esca-BD epidemiology and plant material

The 19 Gamaret plots studied are spread over a perimeter of about 3700 ha of cultivated vineyards in Western Switzerland (Fig. 1). The plots monitored are located in four wine-growing regions with plots located at the shore of the Lake of Geneva (9 plots), of the Lake of Neuchâtel (3 plots), in Chablais (3 plots), all at an altitude of 350 m., and in Valais (4 plots) located in a large alpine valley between 500 m and 700 m altitude.

**Fig. 1.**
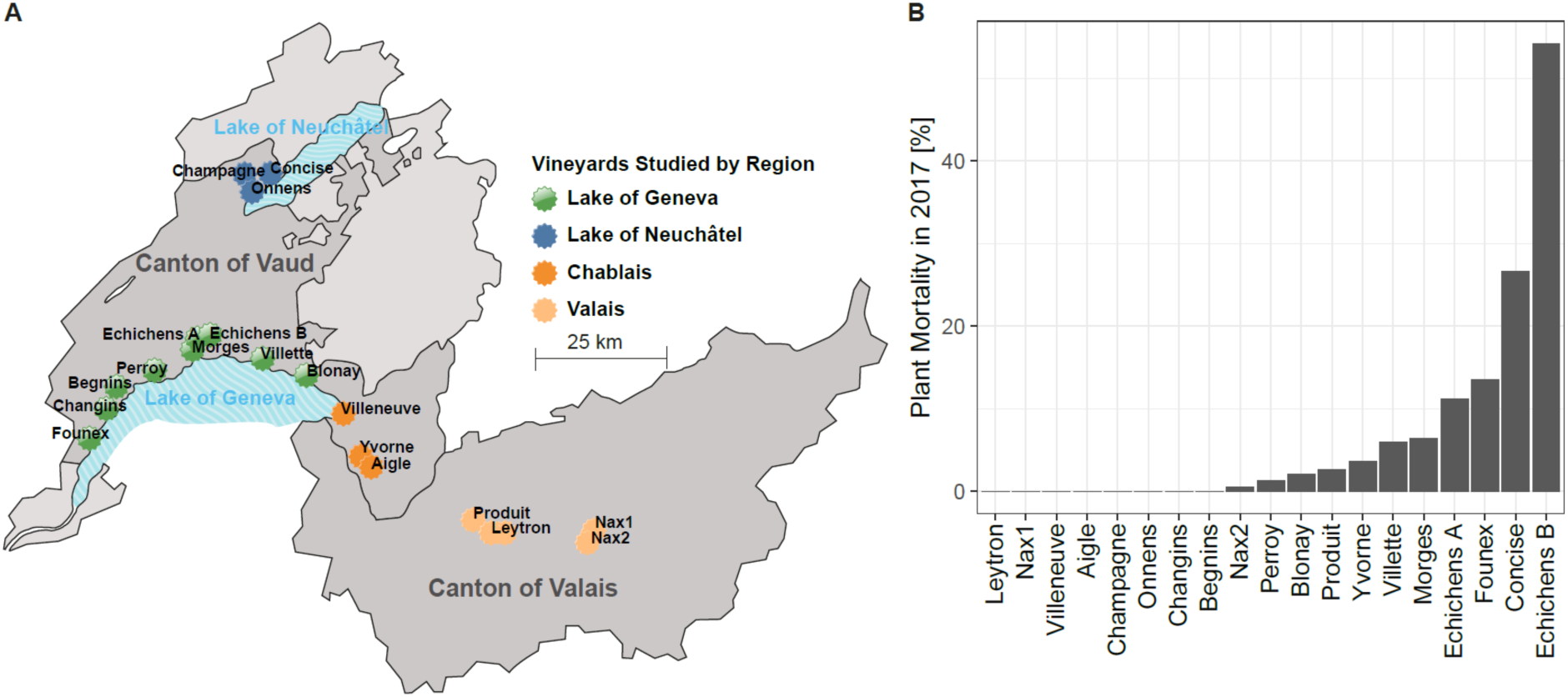
**A** Map of the Gamaret vineyards monitored across western Switzerland. **B** Cumulative replacement rate [% of plants] from vineyard plantation in 2003 through 2017 across the study plot network.

These plots were all planted in 2003 with *Vitis vinifera* L. cv Gamaret, a Swiss red variety developed from a cross between Gamay x Reichensteiner. This vine variety is well established in Switzerland with circa 500 ha (representing 3% of the Swiss vineyard area). Vine plants were all grafted onto 3309C rootstock (*V. riparia x V. rupestris*) with the same vegetal/genetic material coming from the same nursery stock. All vines were trained in the single Guyot system (vertical shoot-positioned foliage) with an average planting density of 7500 ± 500 vines ha^-1^. The soil management included a natural grassing between the rows for all the plots implanted in Vaud (15 plots, Fig. 1. A). On the Valais network (4 plots), chemical weed control was carried out.

The vineyard plots (Fig. 1B) were monitored for GTD leaf symptoms and mortality since their planting (2003) until the beginning of this study (2018). Foliar symptoms (Fig. 2B) in the studied vineyard plots were typical of esca and/or Botryosphaeria dieback (BD). As we did not identify the fungal pathogens associated with the vineyard plots studied, we refrained to refer to a specific GTD in the present paper. Foliar, symptoms are difficult to attribute to esca and/or Botryosphaeria dieback (Luque et al., 2009; Viret & Gindro, 2014.). Consequently, we used esca-BD to qualify the disease plants were suffering from in the vineyard network studied. Cumulated plant mortality data gathered before the experiment (2003-2017) appeared highly variable (Fig. 1B), ranging from 0% to 48 % depending on the vineyard plots. Consequently, Gamaret cultivar seemed to be a good candidate to study the influence of biotic and abiotic factors on the incidence of esca-BD.

**Fig. 2.**
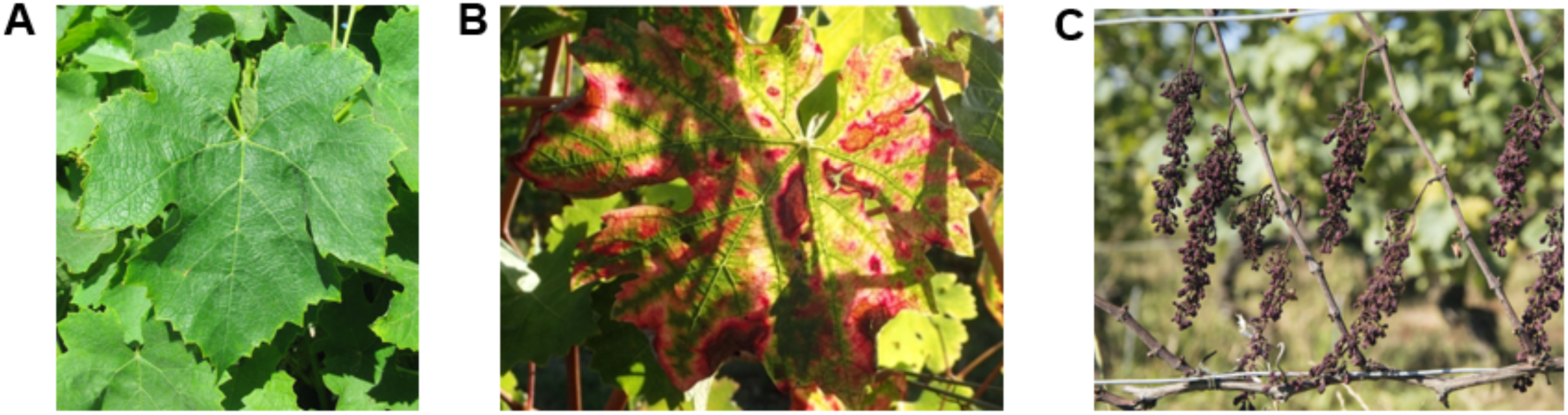
Epidemiological data recorded according to visual symptoms A. Asymptomatic leaves; B. Leaves expressing esca-BD symptoms; C. Apoplectic plant.

For four consecutive experimental years (2018-2021), when esca-BD symptoms were most visible in late August or early September, about 300 vines in each of the studied vineyard plots were examined for the presence of esca-BD symptoms (5700 plants on the whole network). For the epidemiological monitoring, we identified five categories of symptoms (Fig. 2): apparently healthy plants; foliar symptoms taking in account incidence and severity weak foliar symptoms: one symptomatic shoot, medium foliar symptoms: more than a single symptomatic shoot, strong foliar symptoms: whole plant shoots symptomatic), apoplectic symptoms; replacement of the plant after apoplexy. The proportion of affected plants per plot, from year to year, was inferred based on the number of the plants originally planted in 2003 and still present in 2021.

### Climatic data

Meteorological data were collected from the Data portal for teaching and research (IDAweb; https://www.meteoswiss.admin.ch/services-and-publications/service/weather-and-climate-products/data-portal-for-teaching-and-research.html) of the Federal Office of Meteorology and Climatology MeteoSwiss. Climatic data were collected from meteorologic stations for locations representative of each of the four wine-growing regions under study (Fig. 1.A): Lake of Geneva (Pully), Lake of Neuchâtel (Method), Chablais (Aigle) and Valais (Sion). We collected weather data for the four years of the experiment (2018-2021) as well as for previous years (1990-2017) to derive a climate norm for comparison. Meteorological norms (min, max, quartiles and median) were then computed for these four stations.

In order to increase the accuracy of meteorological data, meteorological stations from Agrometeo (http://www.agrometeo.ch) were chosen according to their proximity to the network plots (Table SM 1). This network consists in automatic meteorological stations. Those stations were thus not suitable for the computations of climatic norms but were used for the measurements during the experiment (2018-2021). We gathered from Agrometeo precipitations, temperatures, and evapotranspiration measurements for 11 stations. Secondary variables were computed from the raw data. From the precipitation data we created nine secondary variables (Table SM 2): monthly precipitation; number of days per month with rainfall; number of days per month with >= 0.1mm, >0.3mm, >10mm, >100mm; absolute deviation from monthly median sum; deviation of the number of days per month with precipitation from the monthly average; number of days per month superior and inferior to the median. From the temperature data we created four secondary variables: number of days per month above and below the median; number of days per month with >25°C; number of unusually hot days (maximum temperature ≥ 30 °C) per month, deviation from average (1991-2020 norm). We consider only meteorological data from April to September, period most likely to influence the incidence of esca-BD foliar symptoms and plant mortality.

### Soil types and soil water holding capacity (SWHC)

Around 80% of the vineyard plots studied are alpine moraines. Moraines can be classified into three types (Letessier & Fermond, 2004): bottom moraines with few stones (< 30% coarse elements), stony moraines (30-60% coarse elements), and gravely moraines (> 60% large elements and stones).

For most of the plots (15), a hole of 1.5 m deep x 1 m wide was dug between two rows of vines. For four plots, the hole was dug to a depth of 2 m because the roots were deeper and bedrock not reached. The SWHC was calculated according to (Letessier & Fermond, 2004). A cultural coefficient was applied for each of the soil textures (Baize & Jabiol, 2011).

### Physiological and agronomical monitoring of the vineyard plots

To determine the vine water status, two approaches were used: the measurement of predawn leaf water potential (Ψ_PD_) and analysis of the carbon isotope composition *(δ^13^C)* in berries at harvest. Predawn leaf water potentials (Ψ_PD_) were measured using a pressure chamber (Scholander et al., 1965) between 2 and 5 a.m., in complete darkness, on eight mature, undamaged, and non-senescent leaves centrally placed in the foliage on each location when evapotranspiration was at the minimum. The mean values of eight leaves per plot were used as predawn water potential variable. The level of water stress of each plot was assigned according to (Van Leeuwen et al., 2009). Predawn water potentials were measured at the veraison (BBCH 83-85) stage once a year in August. The stable carbon isotope (*δ^13^C*) composition of the must sugars (200 berries sample by plot) was determined at harvest at the Stable Isotopes Laboratory of the University of Lausanne by elemental analysis-isotope ratio mass spectrometry (EA-IRMS) using a Carlo Erba 1108 elemental analyzer connected to a Thermo Fischer Scientific (Bremen, Germany) DeltaV mass spectrometer. The stable isotope composition was reported as *δ^13^C* values per mille (‰), with deviations of the isotope ratio relative to known standards as follows: \delta = [(*R*_sample_ − *R*_standard_)/*R*_standard_] × 1000, where *R* is the ratio of heavy to light isotopes (^13^C/^12^C). The *R*_standard_ value for *δ^13^C* in Vienna Pee Dee Belemnite limestone is 0.0112372 (Deléens et al., 1994).

At veraison, a foliar analysis was performed to determine the levels of leaf nitrogen, potassium, phosphorous, calcium and magnesium. The samples consisted of 30 leaves gathered in the cluster zone. Leaves with petioles were washed, over-dried, ground and analyzed by Sol Conseils (Gland, Switzerland). Leaf chlorophyll index was measured using an N-tester device (Yara, Nanterre, France) on adult leaves situated at the middle of shoots. In early winter, the total weight of the pruned vine shoots was recorded (30 shoots per vineyards). The second or last shoot of the fruiting cane was chosen for the measurement. Thirty shoots per plot (one per plant) were sampled and cut to a length of 1.0 meter and individual shoot weights determined (g/linear meter).

At harvest, 200 berries per vineyard were randomly selected and weighted. After weighing the berries, the juices extracted from the individual berries were analyzed by the Oenology Laboratory of Agroscope to establish their sugar, pH, malic and tartaric acid contents, and their assimilable nitrogen content Using WinScan® infrared spectroscopy (FOSS NIRSystems, USA).

### Statistical analyses

We used principal component analyses (PCA) to see if meteorological and physiological parameters could have an influence on annual esca incidence and test if these variables could be used to predict disease incidence. These analyses also feature collinearity between meteorological and physiological variables. Three variables reporting for esca-BD incidence were created: the number of apoplectic plants over the four years of the experiment, the sum of all types of foliar symptoms, and the overall symptoms.

We then used two distinct methods to assess the strength of the relationships between our variables. We first have considered Pearson’s correlation coefficient and a recently developed correlation-coefficient index ξ−correlation coefficient (Chatterjee, 2019) that considers any kind of functional relationship. In particular, it has the property of being equal to zero if and only if there is independence. It has been implemented in R in the package xicor (Holmes & Chatterjee, 2021). To screen the explanatory variables that might influence the esca incidence, we fixed some threshold and considered the variables whose correlation with the variables of incidence was higher (in absolute value for the correlation).

## RESULTS

### Esca-BD Epidemiological data across the four years experiment

Looking at the percentage of plant apoplexy and foliar symptoms in the different regions from 2018 to 2021 (Fig. 3) the plots located on the shores of Lake of Geneva, and to a lesser extent the plots located on the shores of Lake of Neuchâtel, expressed more esca-BD symptoms of both types than the other two regions studied, consistently over the years under study.

**Fig. 3.**
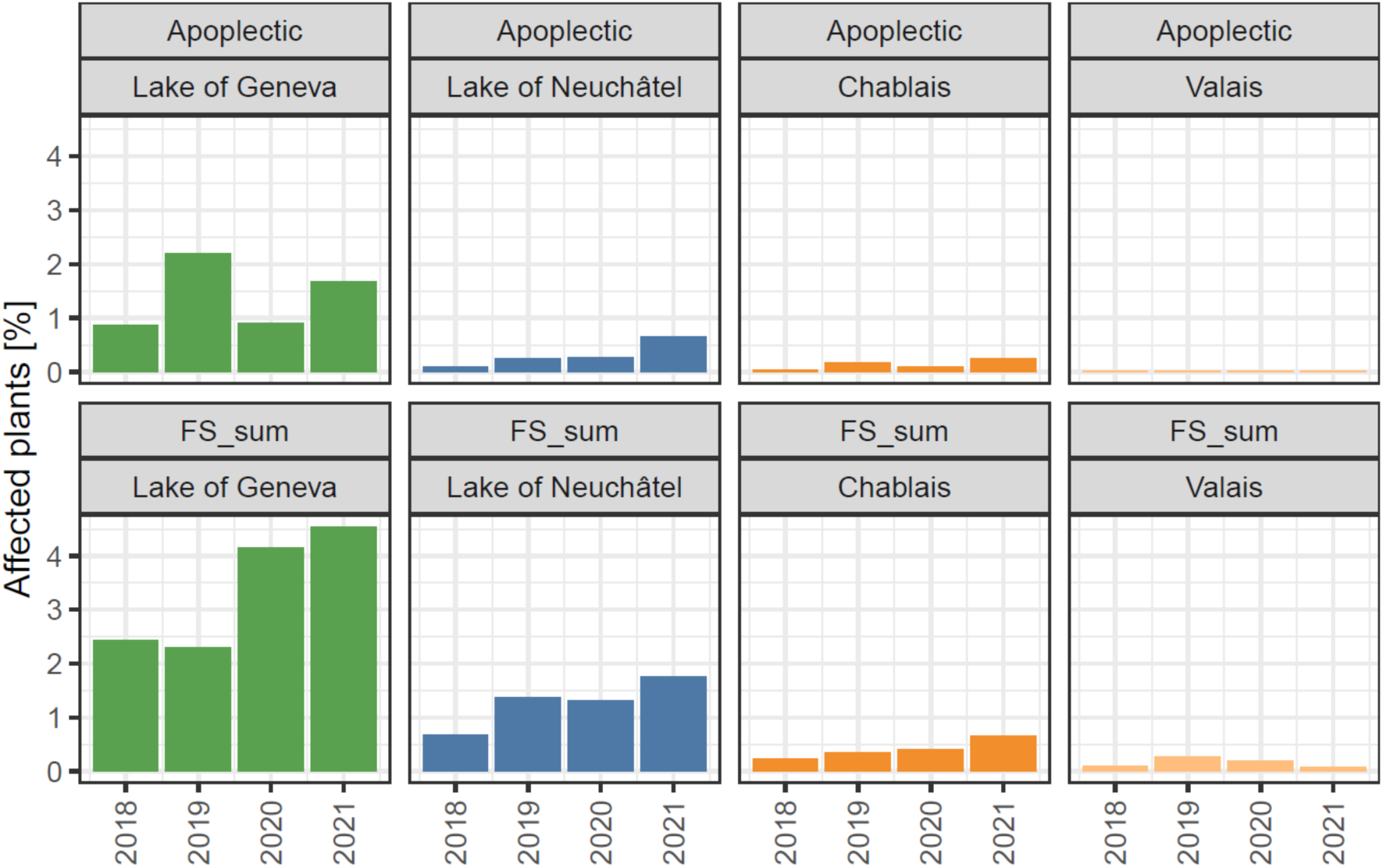
Sum of plants [%] affected by apoplectic symptoms and by sum of foliar symptoms (FS_sum) for the four monitored vintages (2018-2021) by wine-growing region.

In Valais, annual symptoms (apoplexy and foliar symptoms) were close to zero, while in Chablais they were slightly higher but always inferior to 1%. However, apoplexy and foliar symptoms showed a different evolution during the four years of experimentation. The years 2019 and 2021 were the worst for apoplexy in three of the regions (Lake of Geneva, Lake of Neuchâtel and Chablais; Fig. 3), as were the leaf symptoms of esca-BD, but only in Lake of Neuchâtel. In 2020 and 2021, more than 4% of the vines in the Lake of Geneva plots expressed foliar symptoms, compared to just over 2% in 2018 and 2019. Foliar symptoms were generally slightly increasing from year to year in all regions except Valais. Of the four regions studied, Lake of Geneva is clearly the most susceptible to esca-BD, while Valais and Chablais seem much less susceptible.

A closer look at the epidemiological data taking in account individual plots instead of wine-growing regions, indicates that esca-BD also varies greatly between plots in each region (Fig. 4). At Lake of Geneva, Morges appears to be the worst for cumulative symptoms of the disease, ranging from 21% in 2018 and increasing over the years to 56% in 2021. Esca-BD has also higher incidence in Begnins (17-40%), Echichens A (18-27%), and Villette, ranging from 7-18%, than in the other plots of the region considered. However, the incidence of cumulative symptoms did not exceed 10% in Blonay during the four-year experiment. In the Lake of Neuchâtel region, the cumulative incidence of esca-BD also varied. While Onnens expressed the lowest percentage of esca-BD symptoms over the years 2018-2020 (4-11%), Champagne showed slightly more symptoms over the same period (8-12%) and Concise was the most affected plot (10-25%). However, in 2021, the situation was reversed, with Onnens and Champagne both expressing more than 20% of cumulative esca-BD symptoms while Concise expressed less than half (10%) of symptoms than the other two plots. In Chablais, the plot expressing the highest rate of cumulative symptoms of the disease from 2018 to 2021 was Yvorne (with a peak of 14% in 2018), while the plots in Villeneuve and Aigle expressed 1-2% of symptoms in the first two years and a little more thereafter but still less than 5% except in Villeneuve in 2021 with 9% of affected plants. In Valais, Produit was the plot with the most symptoms, but less than 7%, except for the plot in Nax 2 (5% vs. 4% in Produit) in 2020. In Leytron and Nax 1, the incidence of esca-BD was almost zero in all four years. Overall, these results indicate that esca-BD symptoms incidence varies between years and between plots whatever the region considered.

**Fig. 4.**
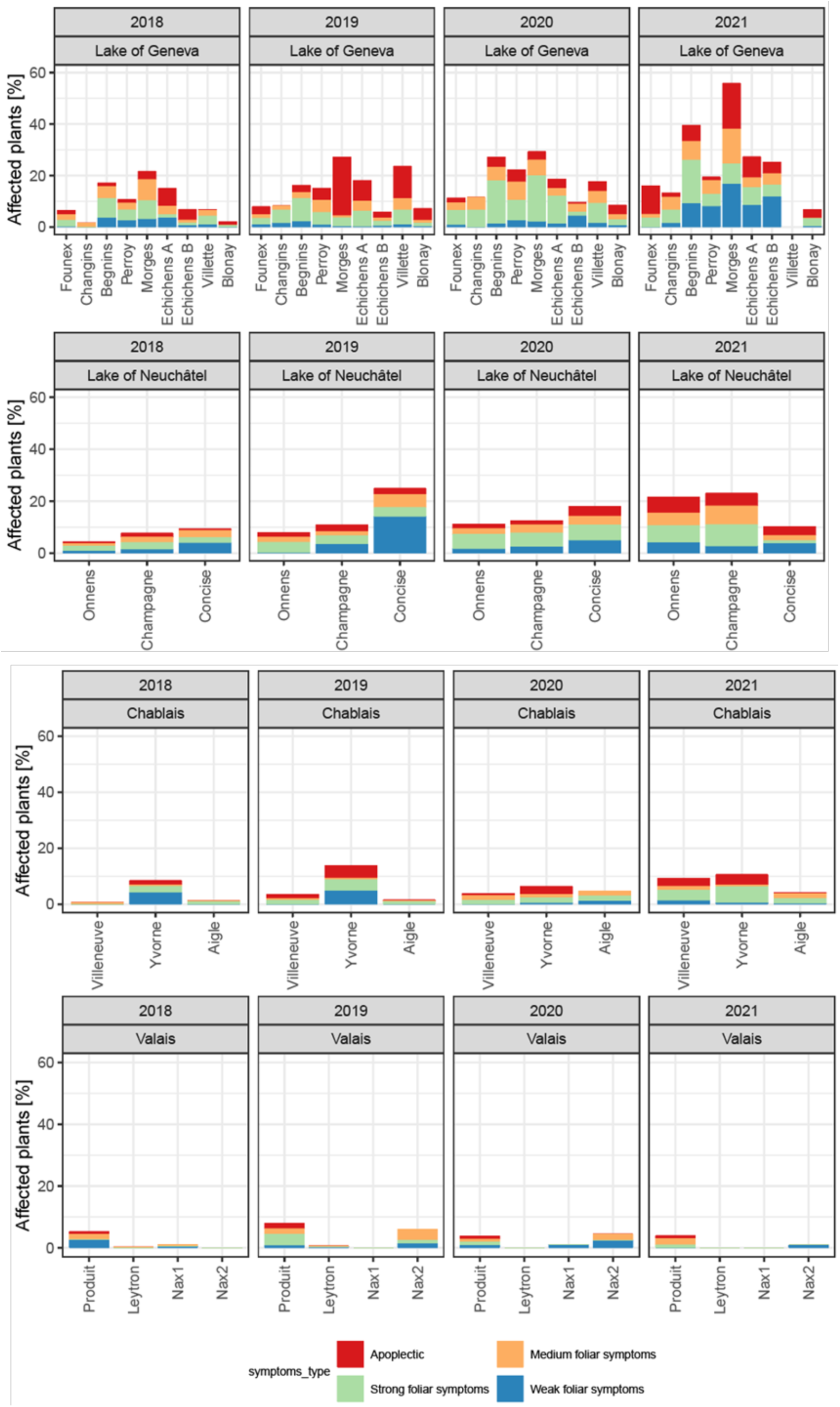
Annual rate of esca-BD symptomatic plants by vineyard plots and by region for weak, medium, strong foliar symptoms and apoplectic plants (2018-2020)

The four-year survey period was marked by unusual precipitation regimes (Fig. 5). While 2018 was drier than the long-term norm, except in Lake of Neuchâtel region from May to July (Fig. 1A), 2019 and 2020 were characterized by an alternance of drier or wetter months than the long-term norm during the six months of grapevine vegetative period considered.

**Fig. 5.**
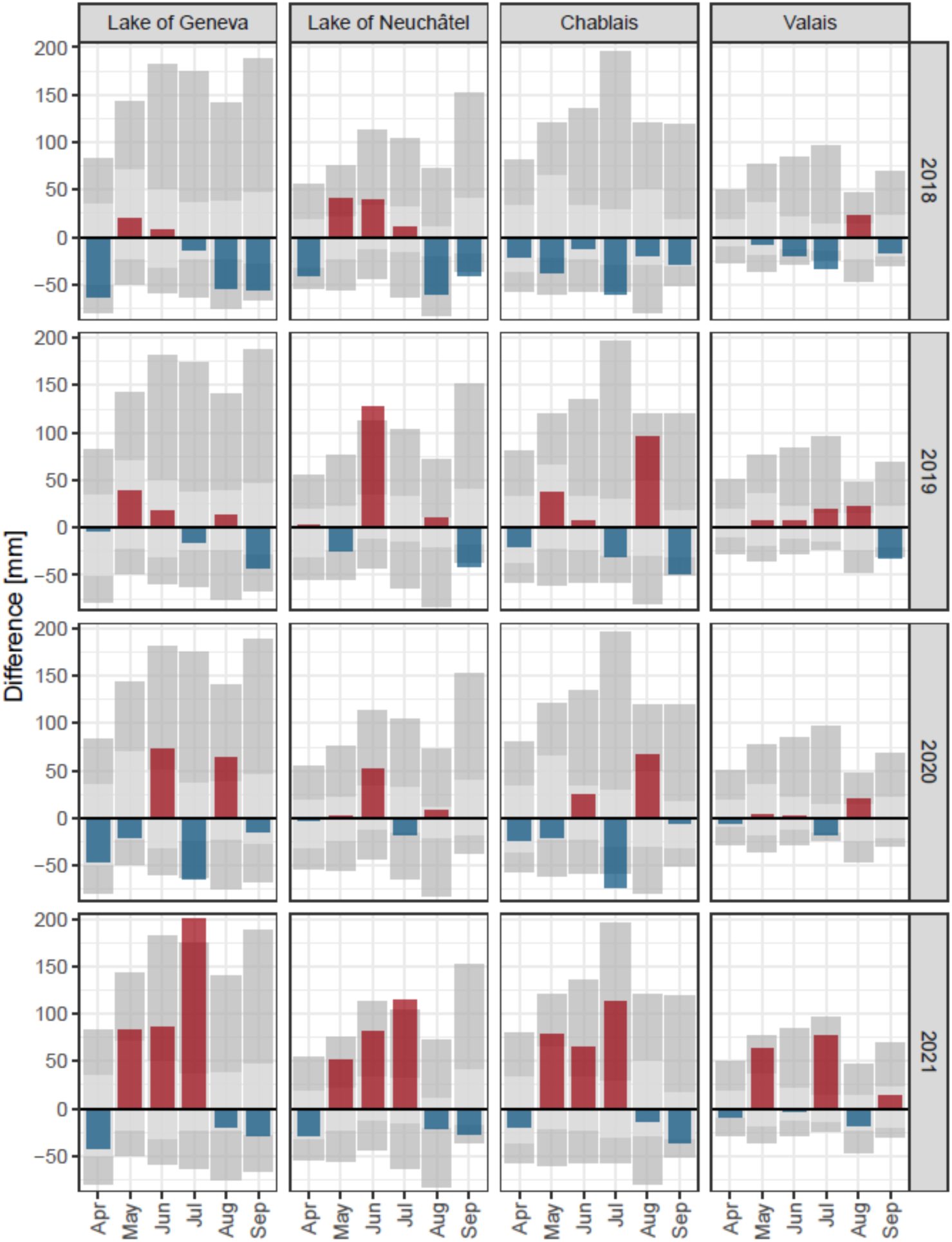
Positive (red) or negative (blue) differences to the long-term median (1990-2017) in monthly spring and summer precipitations (mm) in four wine-growing regions of Western Switzerland from 2018 to 2021. The background rectangles are the minimum and maximum (light grey) and the 25% and 75% percentile (dark grey) of the same data over the period 1990-2017.

Year 2021 encompassed an overall very wet period from May to July, preceded and followed by months drier than the long-term norm, this except for Valais where June was very close to the norm while very rainy in all the other regions. Precipitation differences to the long-term norms were observed in all four wine-growing regions (Fig. 5), however more pronounced in Lake of Geneva, Lake of Neuchâtel and Chablais than in Valais, where except in 2021 in May and July, the precipitation regime remained closer to the norm than in the three other regions.

Temperatures (Fig. 6) were overall higher than the long-term norm during the first three years of the survey period (2018-2020) in the four wine-growing regions, except for May in 2019, which was particularly cold everywhere, and June which was a bit colder than usually in all regions except in Valais. Year 2021 was very peculiar regarding temperatures and highly similar in the four wine-growing regions. This year was characterized by an alternance of cold and hot months in all regions. While April and particularly May were colder than the long-term temperature norm, June appeared hotter than the norm, July and August cooler and September hotter. The month of May appeared the most singular month in 2019 and 2021, with very cold temperature compared to the long-term norm in all four regions. Also, during the four years of the survey period, temperatures were more homogenous across regions (Fig. 6) than the precipitations (Fig. 5) and Valais appeared not different than the other three wine-growing regions regarding temperature fluctuations.

**Fig. 6.**
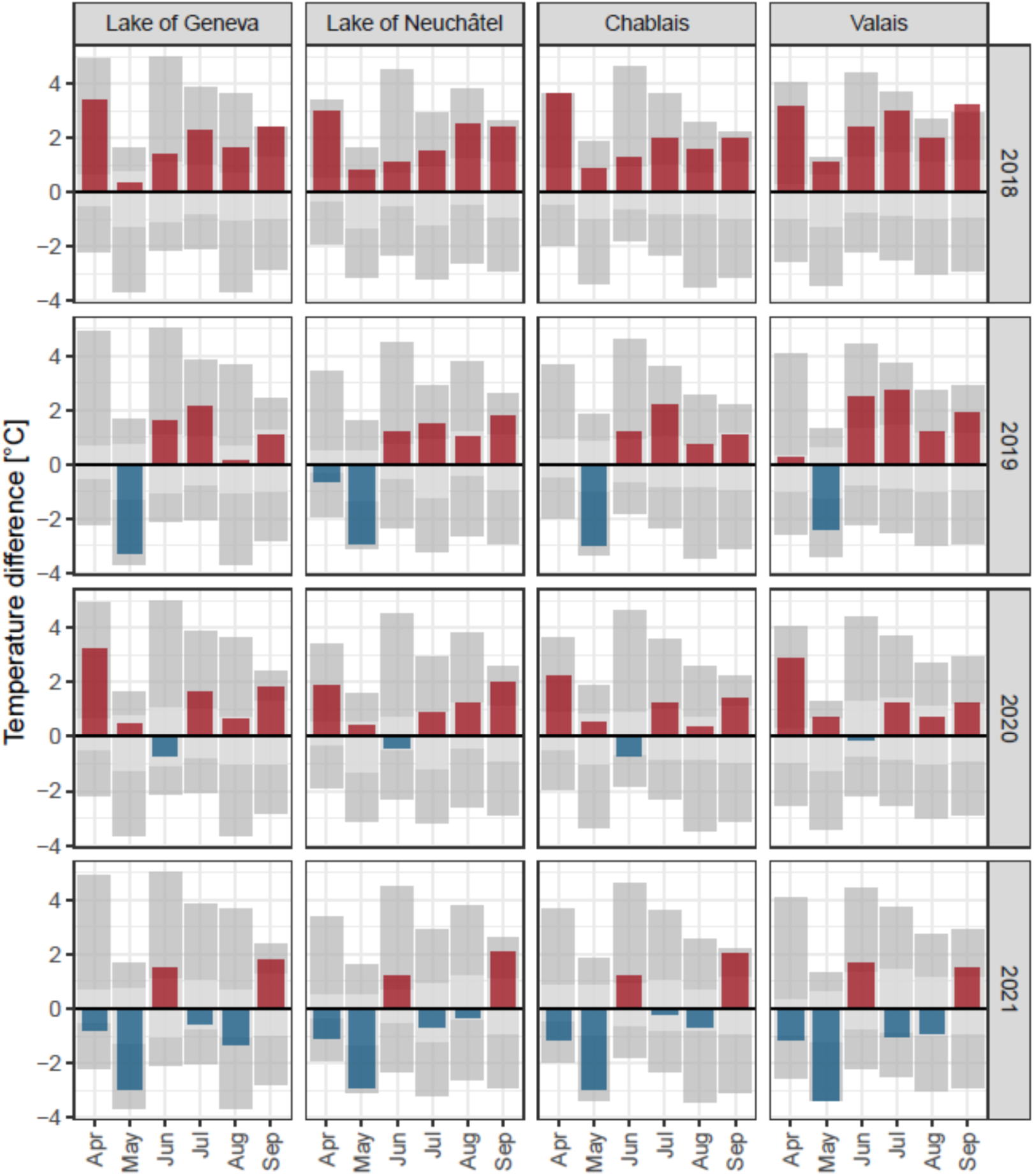
Positive (red) or negative (blue) differences to the long-term median (1990-2017) in monthly spring and summer temperatures (°C) in four wine-growing regions of Western Switzerland from 2018 to 2021. The background rectangles are the minimum and maximum (light grey) and the 25% and 75% percentile (dark grey) of the same data over the period 1990-2017.

### Soil types and soil water holding capacity (SWHC)

The sites studied were roughly grouped by category of SWHC (Table 1).

**Table 1.** Experimental sites in four wine-growing regions (Switzerland), with their soil type managements (% soil cover cropping) and their soil water holding capacity (SWHC).

| Sites | Region | Soil types | % Cover cropping | SWHC (mm) | SWHC category |
| --- | --- | --- | --- | --- | --- |
| Founex | Lake of Geneva | Gravelly moraines | 70 | 155 | Medium |
| Changins | Lake of Geneva | Bottom moraines | 70 | 110 | Medium |
| Begnins | Lake of Geneva | Bottom moraines | 70 | 130 | Medium |
| Perroy | Lake of Geneva | Bottom moraines | 70 | 155 | High |
| Morges | Lake of Geneva | Bottom moraines | 70 | 240 | High |
| Echichens A | Lake of Geneva | Bottom moraines | 70 | 230 | High |
| Echichens B | Lake of Geneva | Bottom moraines | 70 | 230 | High |
| Villette | Lake of Geneva | Marly sandstones | 70 | 250 | High |
| Blonay | Lake of Geneva | Marly sandstones | 70 | 200 | High |
| Onnens | Lake of Neuchâtel | Jurassic sandy stones | 70 | 120 | Medium |
| Champagne | Lake of Neuchâtel | Jurassic sandy stones | 60 | 95 | Low |
| Concise | Lake of Neuchâtel | Jurassic sandy stones | 70 | 180 | High |
| Villeneuve | Chablais | Gravelly moraines | 70 | 90 | Low |
| Yvorne | Chablais | Gravelly moraines | 50 | 100 | Low |
| Aigle | Chablais | Gravelly moraines | 50 | 80 | Low |
| Produit | Valais | Clay schists | 0 | 160 | High |
| Leytron | Valais | Stony moraines | 0 | 100 | Low |
| Nax1 | Valais | Stony moraines | 0 | 90 | Low |
| Nax2 | Valais | Colluvial/stony moraines | 0 | 140 | Medium |

The SWHC was calculated on each plot by taking into account the amount of stones, texture, root colonization and rooting depth (Letessier & Fermond, 2004). The SWHC corresponds to the maximum amount of water in a soil that the vine can extract (Baize & Jabiol, 2011). A quarter of the plots have a SWHC ≤ 100 mm (low), another quarter of the plots have a SWHC between 100 and 150 mm (medium), and half of the plots a SWHC > 150 mm (high).

### Physiological and agronomical monitoring of the vineyard plot

Comparing the different record year, none of the plant physiological indices (Ψ_PD_, δ^13^C, assimilable nitrogen, chlorophyll content of the leaves, weight of the pruning and of the berries) remained stable across the four years experiment (Fig. 7). Apart from water potential (Ψ_PD_), which increased in 2020 for plots with an average SWHC, remained stable for plots with a high SWHC and decreased for plots with a low SWHC (Fig. 7A), all the other biotic factors (Fig. 7B-F) followed the same pattern depending on the vintage and independently of the SWHC. Grapevine plants grown on soils having a low SWHC exhibit lower pre-dawn water potential and pruning weight but a higher rate of assimilable nitrogen and chlorophyll than vineyards planted in soils with medium and high SWHC. For these latter plots, assimilable nitrogen in harvested berries (Fig. 7C) was generally low in 2018 and 2021 (with average values between 60 and 80 mg N/litre), tendency less pronounced for plots with low SWHC, and reflecting a marked nitrogen deficiency in grapes during the first and last year of the experiment. Pruning weight (Fig. 7E) was particularly low in 2020 but only for plots having a low SWHC. Berries weight was high in 2019, this independently of SWHC (Fig. 7F). Consequently, the biotic factors associated with the vineyard plots seem to vary more from one vintage to another than according to the SWHC category.

**Fig. 7.**
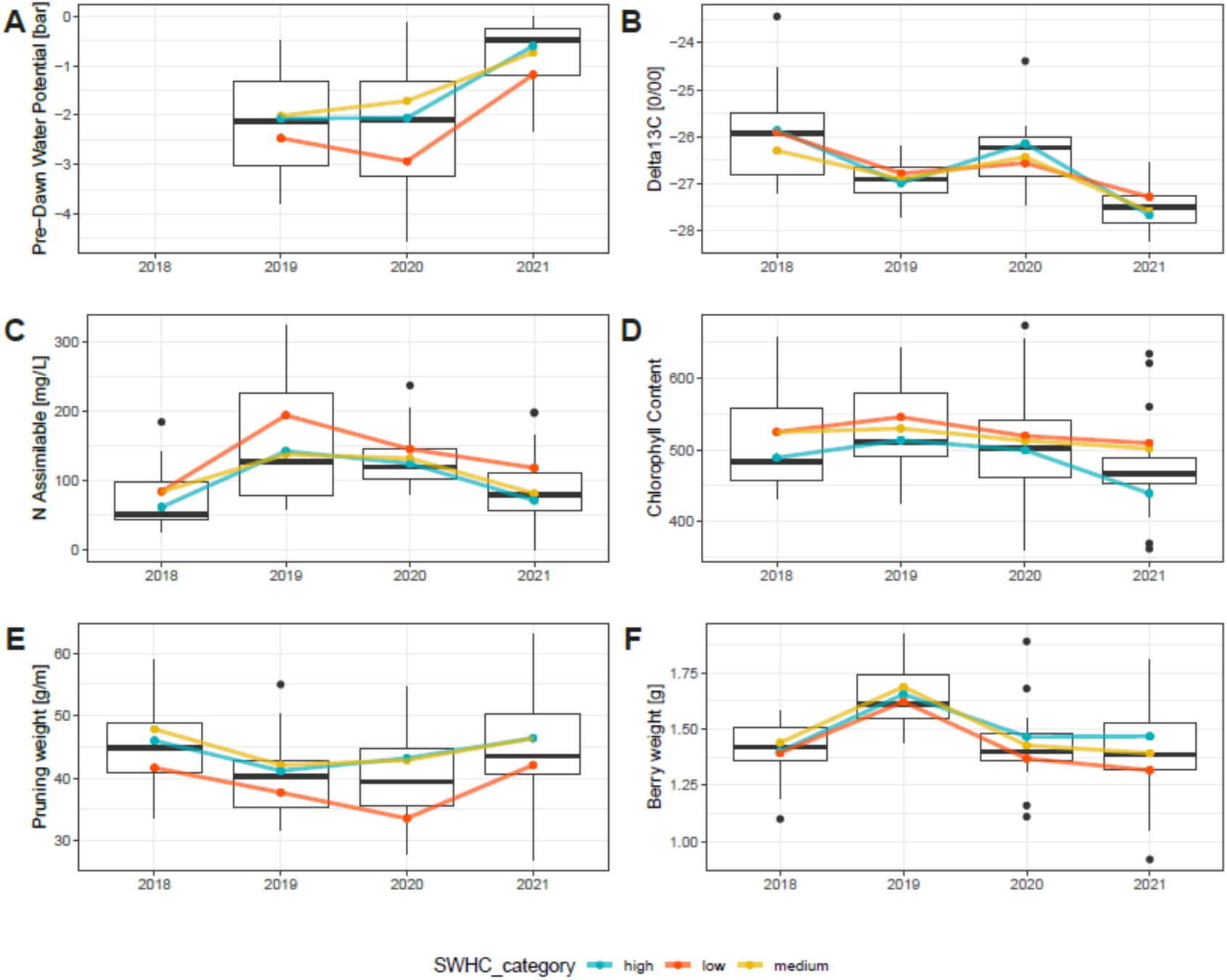
Mean values of measured physiological indexes of plots categorized by SWHC (<100 low [orange], >100-150 medium [yellow], > 150 high [blue]) across four consecutive years (2018-2021) **A** pre-dawn leaf water potential (Ψ_PD_), **B** δ13C, **C** Assimilable nitrogen, **D** Chlorophyll content index (N-tester), **E** Pruning weight, **F** Berries weight.

By grouping the plots by region (Fig. SM1), the physiological variables seem to be more influenced by the regional climates, particularly for the predawn leaf water potential (Fig. S1A), and the nitrogen content (Fig. S1C-D). The Valais plots are atypical according to these indicators, highlighting a regional particularism. The Chablais appears, as the Valais, characterized by a high predawn leaf water potential and a high nitrogen content.

### Multivariate analyses including abiotic and biotic factors

Collinearity between measured abiotic parameters and incidence of the disease with main pattern of variation (Fig. 8) were summarized with a principal component analysis (PCA). First PCA axis explains between 43.8% and 54.8% of the variance depending on vintages and is mainly driven by nitrogen indices, esca-BD incidence indices, cover cropping and SWHC. Second PCA axis explains between 12.9 to 24.9% of the variance and is mainly driven by wood and berries weight and, however to a lower rate, by water stress indices (Delta13C and Base_pot) depending on the vintages.

**Fig. 8.**
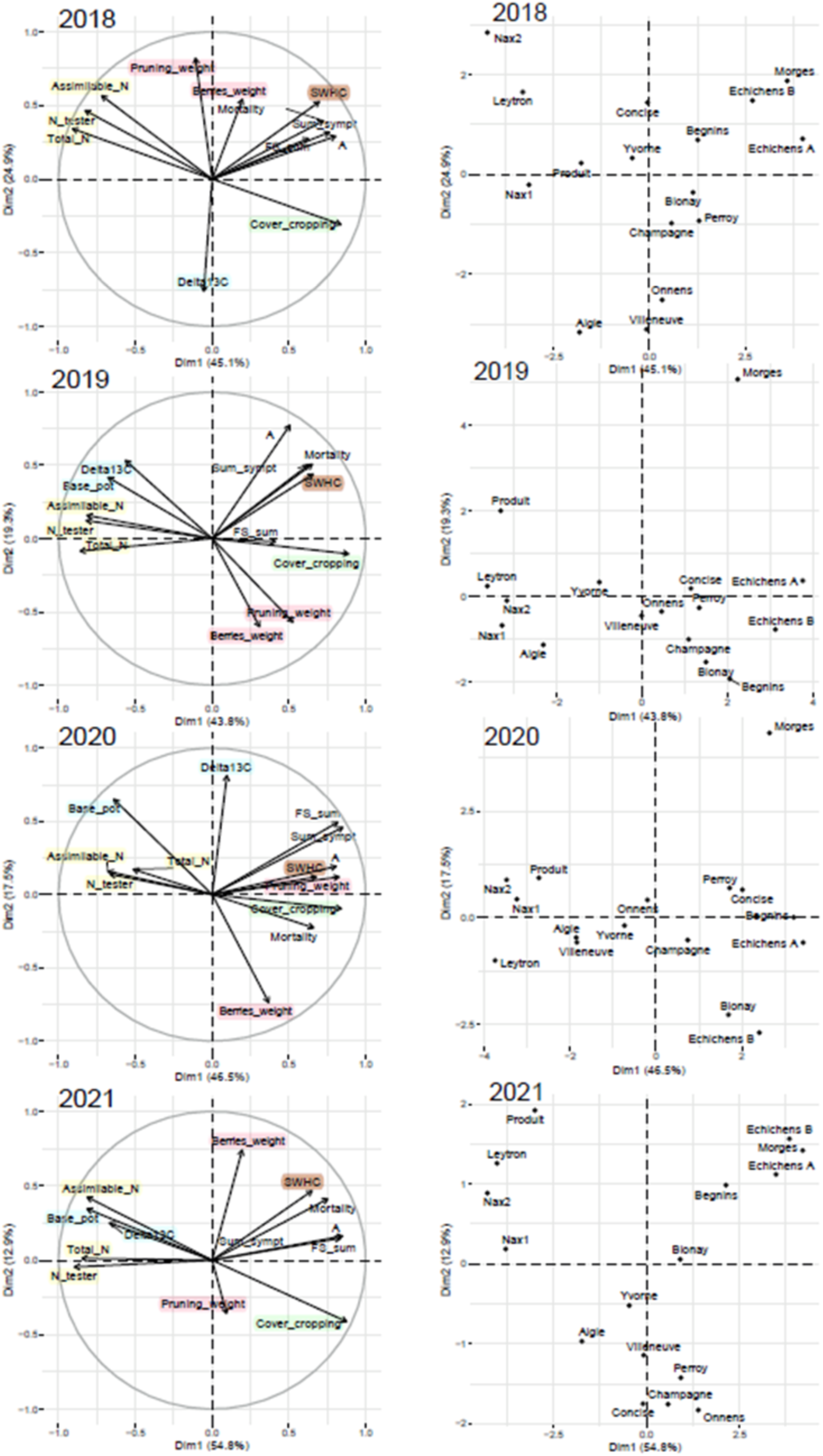
Principal Component Analysis (biplot PCA) based on abiotic and epidemiological variables by vintages 2018-2021 (left) taking in account the 19 plots (right). Color code account for nitrogen content (yellow), vigor (pink), water stress index (blue), cover cropping (green), soil water holding capacity (brown) and epidemiological esca-BD variables (no color) with apoplexy (A), sum of foliar symptoms (FS_sum), all symptoms (A+FS_sum) and mortality.

According to PCA results, SWHC is correlated with the annual incidence esca-BD (arrows point in the same direction). SWHC arrow is closer to plant mortality in 2019 and 2021 and with the sum of foliar symptoms in 2018 and 2020. The variables corresponding to cover cropping, berry weight, and wood weight also point, more than the other biotic factors, in the same direction as the esca-BD incidence indices. This is more pronounced in 2019 and 2021.

Plots located in Valais (Leytron, Nax1, Nax2 and Produit) are clearly discriminated by PCA for the four observed years and are more correlated with nitrogen and water stress indices. These indices were well correlated, especially in 2019 and 2021. These plots are also the least affected by esca-BD (according to annual symptom incidence; Fig. 3). Plots most affected by esca-BD are usually characterized by high SWHC and cover cropping indices. Vigor indices (wood and berry weight) tend to be negatively correlated with high nitrogen indices (more strongly detected in 2019, 2020 and 2021). Cover cropping is always negatively correlated with nitrogen indices.

As SWHC appeared well correlated with plant mortality, the relationship between the total mortality rate since the vines were planted (2003) and the SWHC was tested for the 19 studied vineyard plots. Linear regression (Fig. 9) inferred a positive correlation (R^2^ = 0.6 [P <0.001]) between plant mortality and SWHC. However, while such a correlation is clearly inferred for most of the plots (13 out of 19), SWHC does not explain the plant mortality for six plots, especially for Echichens B, Concise and Villette. However, for the latter, this is probably due to the absence of epidemiological data in 2021, as this vineyard was uprooted in 2020. Nevertheless, the correlation between plant mortality and SWHC suggests that soil water storage capacity is one of the main factors explaining the incidence of esca-BD mortality for the Gamaret cultivar.

**Fig. 9.**
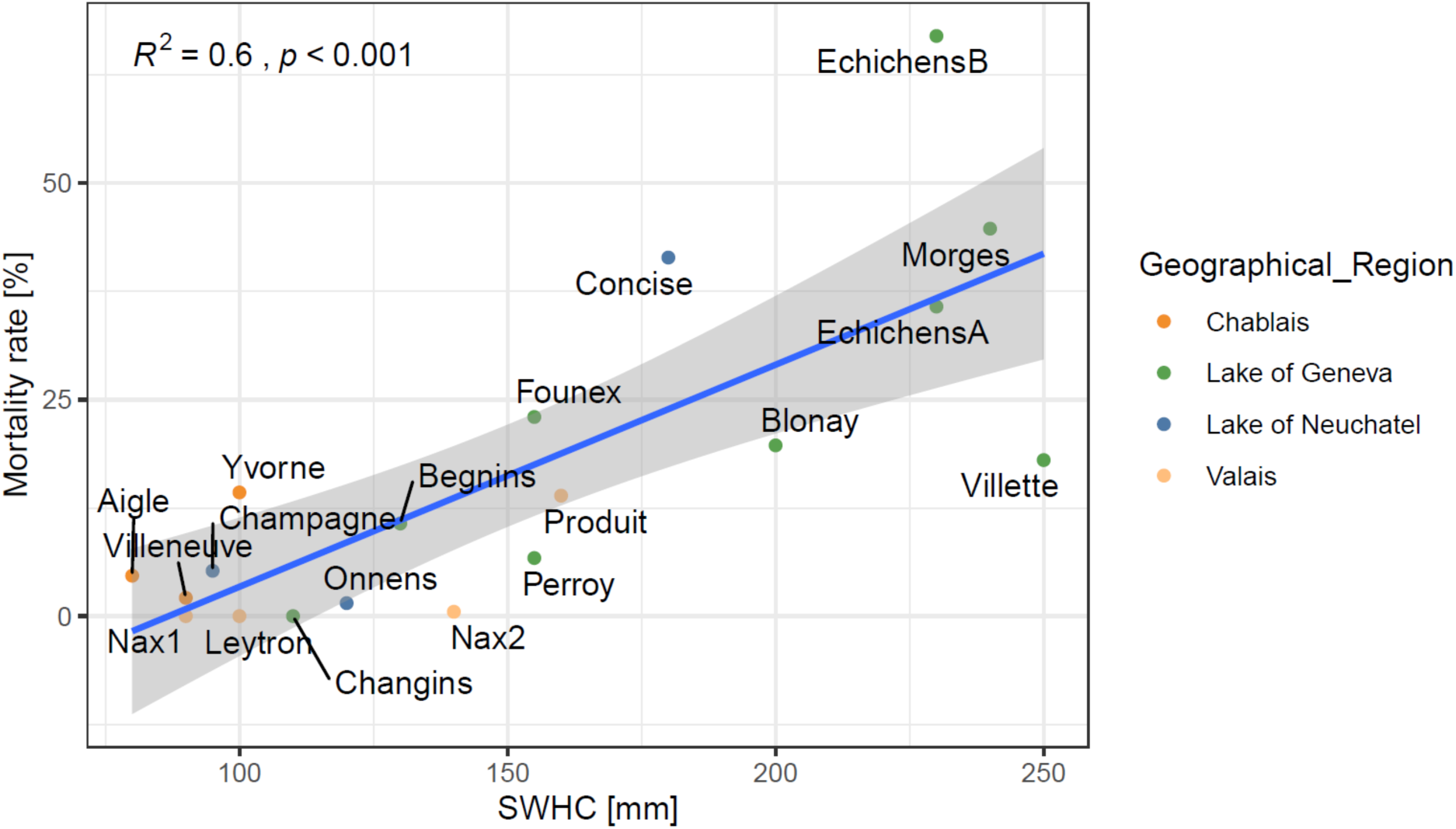
Linear regression between total mortality rate by vineyard plot (from 2003-2021) and soil water holding capacity (SWHC) index for each plot based on Pearson’s correlation coefficient.

According to our correlation test among 70 variables (Table SM 2) with our esca-BD incidence indexes, the variables most correlated are variables accounting for monthly precipitation regimes in May and June, SWHC, cover cropping and chlorophyll content for Pearson’s correlation coefficient (PCC) (Fig. 10). The highest PCC obtained (0.58) describe a positive relation between the sum of symptoms and the rainfall in June (Fig. 10A). PCC also underscore a positive correlation for the precipitation regime observed in June and May between the amount of rainfall and the number of days with rain >10mm in May and June and the difference of precipitation compared to the norm (1991-2017) in June and our esca-BD incidence indices (Fig. 10A). SWHC is also positively correlated with all the esca-BD incidence indices but more strongly with apoplexy. Chlorophyll content was negatively correlated with our esca-BD incidence. This negative relationship between nitrogen and esca-BD incidence was also underscored by the PCA (Fig. 8). Variable most linked with esca-BD incidence indices according to ξ-correlation (XCC) are the rainfall in June and to a lesser extent in July with only apoplexy. Cover cropping and SWHC are also retained as not independent from our esca-BD incidence indices (Fig. 10B).

**Fig. 10.**
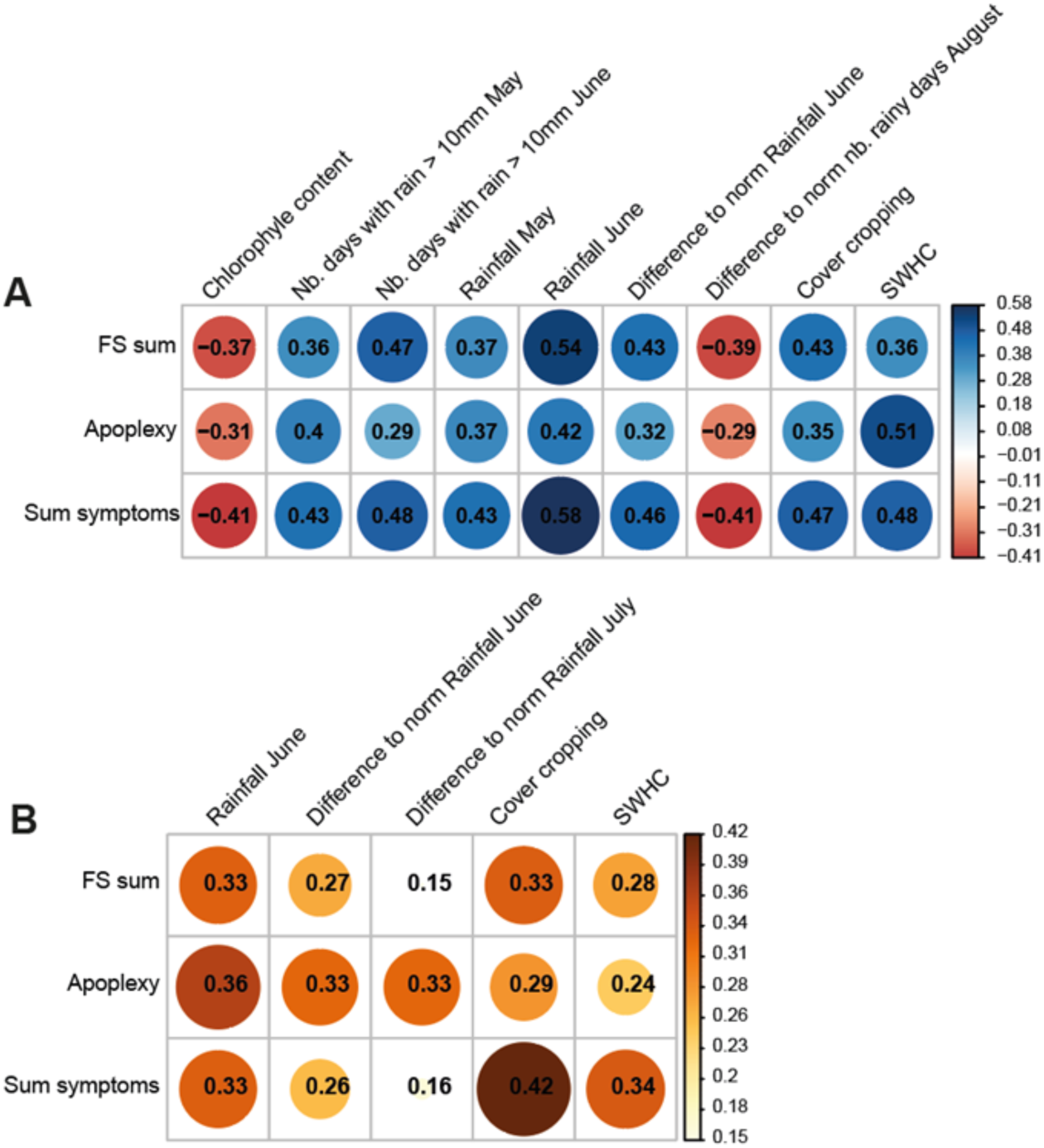
Correlation coefficient matrix with variables most correlated with the esca-BD incidence indices: A (plant with apoplexy), FS_sum (plant with foliar symptoms), Sum_sympt (all plants affected by esca-BD; A+FS_sum). Size of the dot reflects the strength of the correlation. **A** Variables with the highest correlation rate with the esca-BD incidence indices according to the Pearson correlation coefficient (red corresponds to a negative relationship and blue to a positive relationship). **B** Variables most correlated with the esca-BD incidence indices according to Xicor correlation coefficient.

## DISCUSSION

In monitoring prior to this study (2003-2017), Gamaret showed a large variability in susceptibility to esca-BD between plots in the study network. Such a wide range of susceptibility was rarely observed for other grape varieties and already suggested an impact of soil-climate factors on disease expression. By relating the incidence of esca-BD to regional climatic conditions, our results show that in a region like Valais, with a drier climate and with June and July hotter compared to other wine-growing regions in Western Switzerland, the incidence of the disease is almost null. Also, the year 2018 was characterized by an abnormally dry and hot vegetation period compared to the other three years. This vintage was also the one in which esca-BD symptom expression was the lowest in the four viticultural regions. Grapevines are often grown at the limit of water stress, providing less water than the plant can transpire (Castellarin et al., 2007; Chaves et al., 2010). This irrigation strategy aims to improve grape quality and reduce water consumption but was reported to increase the sensitivity of plants to heat waves (Edwards et al. 2011). The present study suggests that heat and water stress may not be as deleterious to viticulture as generally assumed (Fernandez et al., 2023; Fischer & Peighami-Ashnaei, 2019; Songy et al., 2019). Our results agree with those of Bortolami, Gambetta, et al. (2021) who showed that water-stressed grapevines do not express esca-BD symptoms, while watered plant express them. Our results run counter to the idea that heat combined with drought exacerbates plant damage (Pandey et al., 2015). As Bortolami et al. (2021) pointed out, the susceptibility of a given grape cultivar to esca-BD does not seem to be altered in the long term by drought, an assumption verified in this study for Valais. Gamaret shows a susceptibility to the disease that is primarily regional, but that varies from one vintage to another (Fig. 3).

Over the four years that these vineyards were monitored, temperatures in each region for a given vintage were comparable, with the same monthly deviations from the long-term norm. The differences in esca-BD incidence observed between wine-growing regions thus appear to be due to annual precipitation more than to temperature patterns. Previous studies mentioned that cool, rainy summers favored the expression of esca-BD foliar symptoms, whereas hot and dry summers favored apoplexy (Surico et al., 2000; Marchi et al. 2006). We did not observe this trend. The year 2018 was particularly hot and dry during the summer and was the year with the lowest number of apoplectic events and foliar symptom rate. The year 2021 does not fit this trend either, as it was the year with the most rainfall and the highest number of foliar symptoms and apoplectic events in three of the four viticultural regions monitored. Apoplectic events were most numerous in 2019 and 2021 in three of the wine regions. Both years were characterized by a particularly rainy and cold May, followed by a hot (2019) or rainy and hot (2021) summer. These observations are rather in agreement with Calzarano et al. (2018) who showed that temperature and precipitation from May to July, or even only in July (in Italy), are key factors for the expression of esca-BD. In our four years of monitoring, the month that has an impact on plant apoplexy is May, if it is cold and rainy, followed by a hot summer (2019) or alternating periods during which temperatures vary when the summer is particularly rainy (2021).

The Gamaret plants studied here were grafted onto rootstock 3309C reported to have low drought tolerance (Verslype et al., 2023). After our results, this cultivar appears to be less sensitive to drought than expected, particularly in 2018, the driest year of the monitoring, and more generally in the Valais region. Kondouras et al. (2006) reported that rootstock 3309C may allow scions to maintain higher stomatal conductance and water uptake under water deficit conditions. In addition, de Souza et al. (2022) observed that plants grafted to this rootstock had the slowest increase in water deficit throughout the growing season which is thought to be due its larger and deeper root system. In the *Catalogue des vignes cultivées en France* (http://plantgrape.plantnet-project.org), rootstock 3309C is described as particularly sensitive to water stress when it occurs suddenly during the growing season and as having a poor adaptation to excess water. This could also explain why apoplexy cases were more frequent in 2019 and 2021, years characterized by a sudden change in climatic conditions (between May and June in 2019) or by a particularly rainy summer accompanied by alternating cool and hot periods (2021). These observations agree with previous studies (Surico et al., 2000) that suggested that apoplexy was related to alternating dry, hot periods and wet, cool periods, conditions leading to high leaf and canopy production and high evapotranspiration that could lead to disruption of sap flow due to gas embolism. Bortolami et al. (2022) found that hydraulic failure and occlusion of vessels by tyloses and gels induced at a distance by esca-BD are not mutually exclusive and that occlusions lead to hydraulic misfunction in the veins of esca-BD symptomatic leaves.

The variability of esca-BD expression observed between wine-growing regions was also observed between plots within the same wine-growing region (Fig. 4). For example, the cumulative expression of esca-BD symptoms in the Lake Geneva region varied from 21-56% in Morges during the four years of monitoring, whereas in Blonay it was less than 10%. This intra-regional variability was also observed in the Chablais region, with the plot in Yvorne showing more symptoms than the plots in Aigle or Villeneuve during the four years of monitoring. In Valais, the plot in Produit expressed more esca-BD symptoms than the other two plots in this region during the four years of monitoring. The fact that the expression of esca-BD is variable between plots in the same wine-growing region, and thus subject to similar climatic conditions, suggests that rainfall and temperature only partially explain the incidence of the disease. These results confirm the idea, put forward by several authors (Marchi et al., 2006; Mugnai et al., 1999; Surico et al., 2000), that other abiotic factors than climate are involved in esca-BD expression.

The abiotic factor, apart from climate, most often suggested to have an impact on esca-BD expression is the soil water holding capacity (Calvo-Garrido et al., 2021; Calzarano et al., 2018; Graniti et al., 2000; Lecomte et al., 2009; Sosnowski et al., 2011; Van Niekerk et al., 2011). Grouping the plots into three categories based on their SWHC (Table 1) and testing its relationship with some physiological and agronomic factors, results show that biotic factors vary more with vintage than according to plot SWHC category (Fig. 7). Although water stress was moderate during the four years of the experiment, the physiological behavior of Gamaret seems to be more sensitive to sudden changes in climatic conditions than to the accessibility of water, probably in relation with the root capacity of rootstock 3909C to access water deep in the soil during dry periods (de Souza et al., 2022). Soil management seems to play a role in the accumulation of nitrogen in the grapes. The Valais plots, the only ones in the network that are not weeded, have higher levels of available nitrogen in the grapes than the plots in the other three wine regions, regardless of the vintage (Fig. SM 1). Cover cropping is mainly grass in these three wine regions. These results are consistent with Abad et al. (2021) who found that cover crops generally do not compete with grapevines for nitrogen, except for grasses. Also, foliar symptoms are less expressed in Valais than in the other wine regions considered in this study (Fig. 3–4), which contradicts the hypothesis that high nitrogen levels increase the severity of foliar symptoms (Lemmens et al., 2004). It also contradicts the idea that dense cultures as vineyards (Gramaje et al., 2018), well supplied with nitrogen, may favor pathogen load (Liu et al., 2017). As suggested by Sun et al. (2020), while nitrogen appears to affect physical defenses and the production of antimicrobial phytoalexins, it also appears to stimulate the production of enzymes and proteins related to systemic resistance. These authors also suggest that, although nitrogen is known to play an important role in plant defense, its impact on pathogen virulence and plant resistance remains understudied.

To determine which biotic and abiotic factors could explain the incidence and severity of esca-BD symptoms, the potential correlation of these factors with the epidemiological data was tested. PCA results (Fig. 8) suggest that esca-BD symptoms are, among all tested factors and across all vintages, most correlated with SWHC, especially variable accounting for plant mortality in 2019 and 2021. Water deficit related to soils with low SWHC has often been suggested as a source of stress for grapevines (Lanari et al., 2015; Lovisolo et al., 2010) and the impact of drought on esca expression often studied (Edwards, Pascoe, et al., 2007; Edwards, Salib, et al., 2007; Fischer & Kassemeyer, 2012; Luque et al., 2009; Ramegowda & Senthil-Kumar, 2015; Surico et al., 2000, 2006). These studies suggested that in the presence of esca pathogens, drought favors the development of disease symptoms. However, Bortolami, Gambetta et al. (2021) found that water stress inhibits esca symptoms expression. This last hypothesis seems to be verified in our study, especially in Valais, which has drier summers than the other regions and where the expression of esca-BD is the lowest (Fig. 3–4,8). But it is also possible that Valais vineyards are less infected by esca-BD fungal pathogens than the other wine regions considered. A study of the fungal community associated with the studied vineyards will allow to answer that question.

The cost of plant replacement is high and is one of the major concerns of winemakers (Bertsch et al., 2013; Gramaje et al., 2018; Hofstetter et al., 2012). Having identified SWHC as one of the main factors related with plant mortality, we performed a linear regression analysis to establish the relationship between plant mortality rate per plot and SWHC (Fig. 9). Regression analyses identified SWHC as clearly correlated with esca-BDA plant mortality. This trend is observed at regional and at the intraregional level. In Valais, the impact of SWHC on plant mortality appears much less pronounced than in the other wine-growing regions. SWHC is less impacting plant mortality when regional climatic conditions are hot and dry. At intraregional level, Produit, the only plot in Valais with a high SWHC has the highest mortality rate in this wine region. The same tendency is observed for Concise (Lake Neuchâtel) and for the three plots with an average SWHC compared to the other plots located at the edge of Lake Geneva, which have a high SWHC. The fact that the three plots of the Chablais have all a low SWHC may also explain the low rate of esca-BD symptoms and mortality expressed in that region.

Although the correlation coefficient of the regression analysis remains relatively low to assess strong relationships, the influence of water availability on esca prevalence, a hypothesis already advanced by several studies (Andreini et al., 2014; Marchi et al., 2006; Surico et al., 2000), is strongly suggested here to be the limiting factor for plant mortality. Regression analyses allowed us to sort out among 70 abiotic variables that water availability in June given by the amount of rainfall, the number of days where it rained more than 10 mm and the difference of precipitation compared to the average (1990-2017), were the factors the most correlated with esca-BD incidence. The precipitation regime in June would likely be the one that, among the variables considered, influences the most the prevalence of esca-BD for a given year. As precipitation regime in May and August were also retained for specific monitored years, we can consider that the precipitation regime during the spring and early summer where the vegetation growth is the greatest (Keller, 2020) will be determinant for esca-BD prevalence. These results, confirm those of Calzarano et al. (2018) who also showed that climatic conditions from May to July, particularly July in Italy, are driving esca-BD prevalence. Although the role of water is increasingly established, the variability in the incidence of esca within a plot, with some plants being affected earlier than others, is not yet clear. The precise role of water availability or restriction in the expression of esca symptoms has yet to be clarified.

Fischer and Peighami-Ashnaei (2019) suggested that environmental factors were driving the prevalence of esca-BD, primarily due to the time lag (often several years) between pathogenic fungal infection of esca-BD and symptom expression (Di Marco & Osti, 2008). This hypothesis, already suggested by previous studies (Marchi et al., 2006; Mugnai et al., 1999; Surico et al., 2000) seems to be verified in our four-year vineyard network monitoring. Both climatic and soil factors (SWHC) seem to have a strong impact on disease expression. Climate seems to be responsible for the differential expression of leaf symptoms of esca-BD and apoplexy, both at the regional and intra-regional level. Apoplexy also appears to be expressed more than foliar symptoms under specific climatic conditions, when cold and hot periods, deviating from the long-term norm, alternate between May and July. This study identified for the first time the water retention capacity of the soil as a factor limiting plant mortality, a result that remains to be confirmed by studies on other grape varieties susceptible to esca-BD.

Although our four years of vineyard monitoring were characterized by particular climatic conditions compared to the long-term norm, esca-BD symptoms and plant mortality remained low, except for some of the plots of the Lake of Geneva region, where cumulative symptoms reached up to 60% (Morges) and mortality up to 58% (EchichensB). This systemic approach allowed the identification of the combination of factors responsible for the expression of the different types of symptoms of esca-BD in Gamaret. In terms of recommendations to winegrowers, an esca-BD-susceptible cultivar such as Gamaret seems to be better adapted to soils with low water retention capacity and to dry and hot climatic conditions, with the rootstock 3309 providing the scions with a lower sensitivity to drought. Adherence to these recommendations will allow for at least partial control of esca-BD for a susceptible grape variety such as Gamaret, and probably for other susceptible grape varieties, as no plant protection product has proven effective against this disease since the ban on sodium arsenite (Songy et al., 2019).

Climate change and temperature elevation are known to increase the number and severity of fungal infections (Fischer & Kassemeyer, 2012; Nnadi & Carter, 2021). Esca-BD, and more generally all GTD, are considered emerging fungal diseases related to climate change and to human activity (Bertsch et al., 2009; Gramaje et al., 2018). Symptoms similar to those of esca-BD have been observed and described since Antiquity (Mugnai et al., 1999), but this disease has only become a concern in recent decades. There are several reasons why a disease like esca-BD may emerge (Garrett et al., 2021). First, pathogen populations may have change and/or new species introduced. Since the late nineteenth century, most grapevine plants are grafted with European scions and American rootstocks, the latter being resistant to phylloxera (Carton et al., 2007). Young plants are traded worldwide and are a source of newly introduced species everywhere vines are cultivated (Gramaje et al., 2018). For example, *Eutypa lata* (Pers.) Tul. & C. Tul., species responsible for Eutypa dieback, is suggested to have been introduced several times in North America by importation of infected material (Rolshausen et al., 2014; Travadon et al., 2012). Second, host populations may have change. New grapevine cultivars are constantly produced, most of them selected for their resistance to diseases like downy and powdery mildews, respectively *Plasmopara viticola* (Berk. & M.A. Curtis) Berl. & De Toni. and *Erysiphe necator* Schwein., but not to GTD. Third, cultural practices may have changed which is the case in viticulture with the introduction of mechanical pruning which has been reported to increase the number of wounds and being less respectful of sap flow than manual pruning (Gramaje et al., 2018; Lecomte et al., 2018). Pruning wounds are the main route for the entrance for GTD associated fungi (Rosace et al., 2023). Consequently, other reasons than climate change may explain esca-BD emergence.

Also according to Garret et al. (2021), the arguments for attributing a significant role to climate change in disease emergence are the ubiquitous presence of the pathogen(s), the absence of changes in pathogen and host populations that might alter resistance dynamics, the absence of changes in cultural practices, peaks in disease expression that should correspond to the climatic demand of the pathogen (but this assessment is only possible when the climatic demand of the pathogen is well known), and long-term monitoring of the disease pattern to derive a convincing trend in the relationship between climatic conditions and disease incidence. Having used a systemic approach, our study fits most of the criteria to attribute an impact to climate change on esca-BD expression. Esca-BD disease, based on foliar symptoms, is present in all the studied vineyard plots. As we used plants grafted in a unique nursery, with genetically homogenous material all planted the same year, a change in host populations is to discard but cannot be excluded for fungal pathogen populations. Cultural practices were the same for all the plots, except for cover cropping in Valais. The four years of intense monitoring of the vineyard network have inferred a correlation between climatic conditions and disease expression rate.

The systemic approach used allowed to identify the pedoclimatic factors impacting disease expression and can serve as a model to identify the environmental factors most impacting other esca-BD susceptible grapevine cultivars to esca-BD and other GTD associated diseases. Such approach can also be useful to identify the abiotic factors accounting for fungal diseases of other cultivated woody plants and for successful reforestation for which climate and soil characteristics have been shown to be important (Baird & Pope, 2022; Hermoso et al., 2021).

## Acknowledgements

We are grateful to the members of the mycology research group of Agroscope (N. Lecoultre, E. Michellod, A.-L. Fabre, P.-H. Dubuis, A. Melgar, S. Schnee, E. Remolif) for their assistance for the epidemiological monitoring of the vineyards. This work was funded by a research grant from the Canton de Vaud to K. Gindro. Conflict of Interest

## Supplementary material

**Table SM 1.**
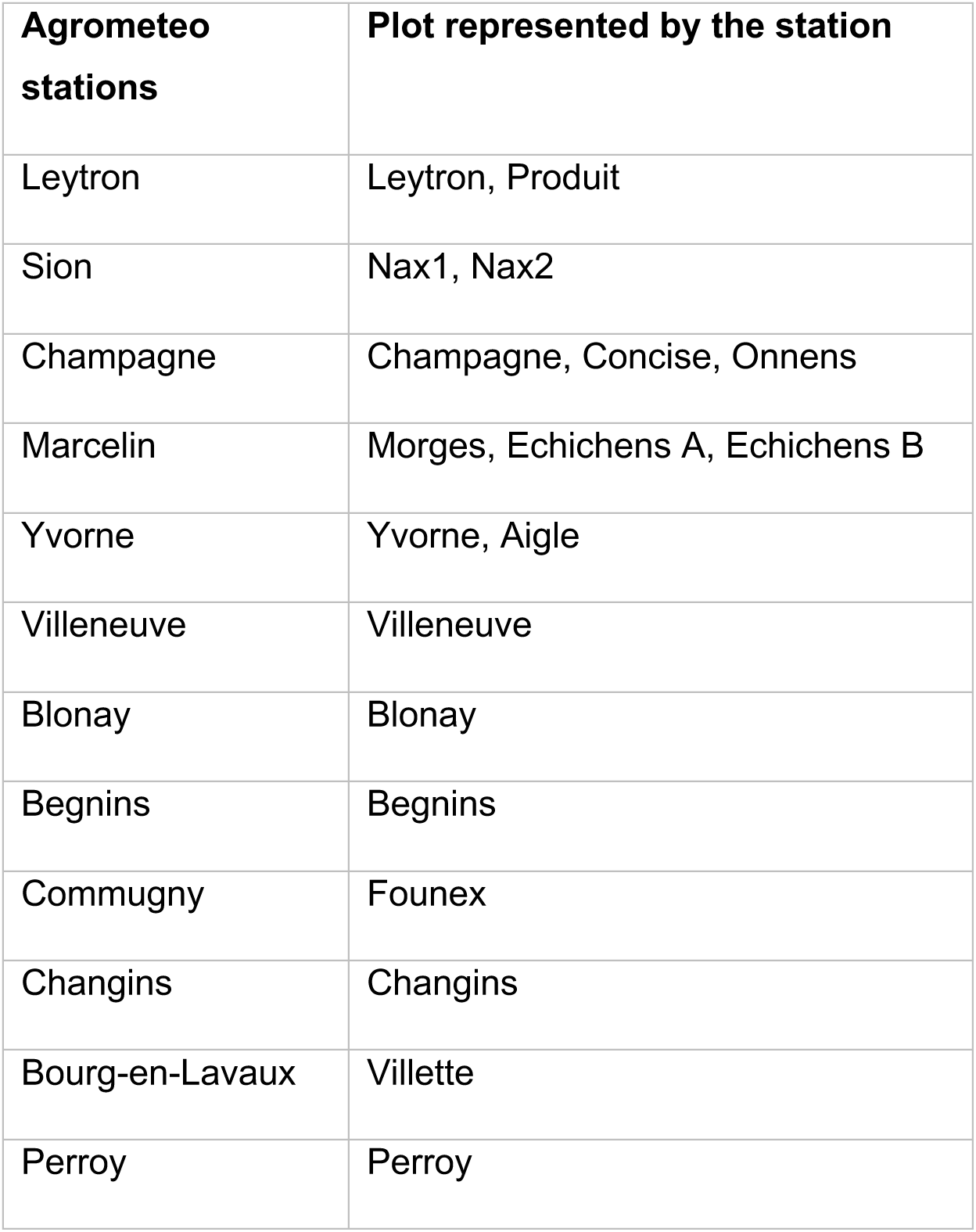
Meteorological station selected for the 19 study sites.

| **Agrometeo stations** | **Plot represented by the station** |
| --- | --- |
| Leytron | Leytron, Produit |
| Sion | Nax1, Nax2 |
| Champagne | Champagne, Concise, Onnens |
| Marcelin | Morges, Echichens A, Echichens B |
| Yvorne | Yvorne, Aigle |
| Villeneuve | Villeneuve |
| Blonay | Blonay |
| Begnins | Begnins |
| Commugny | Founex |
| Changins | Changins |
| Bourg-en-Lavaux | Villette |
| Perroy | Perroy |

**Table SM 2.**
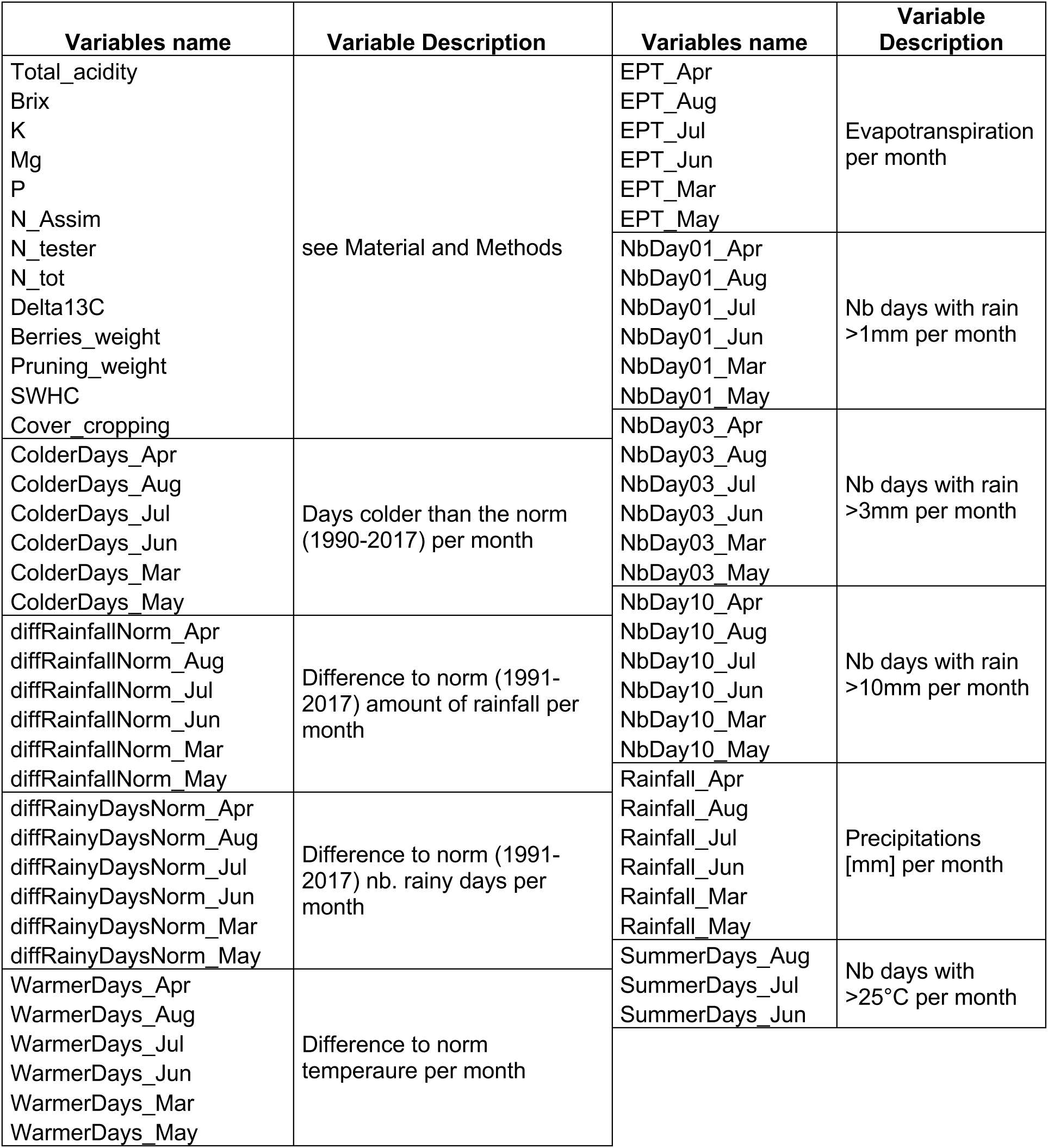
Biotic and abiotic variables considered to establish Pearson and Xicor correlation coefficients with esca-BD epidemiological monitoring data on the entire plot network.

**Figure SM 1.**
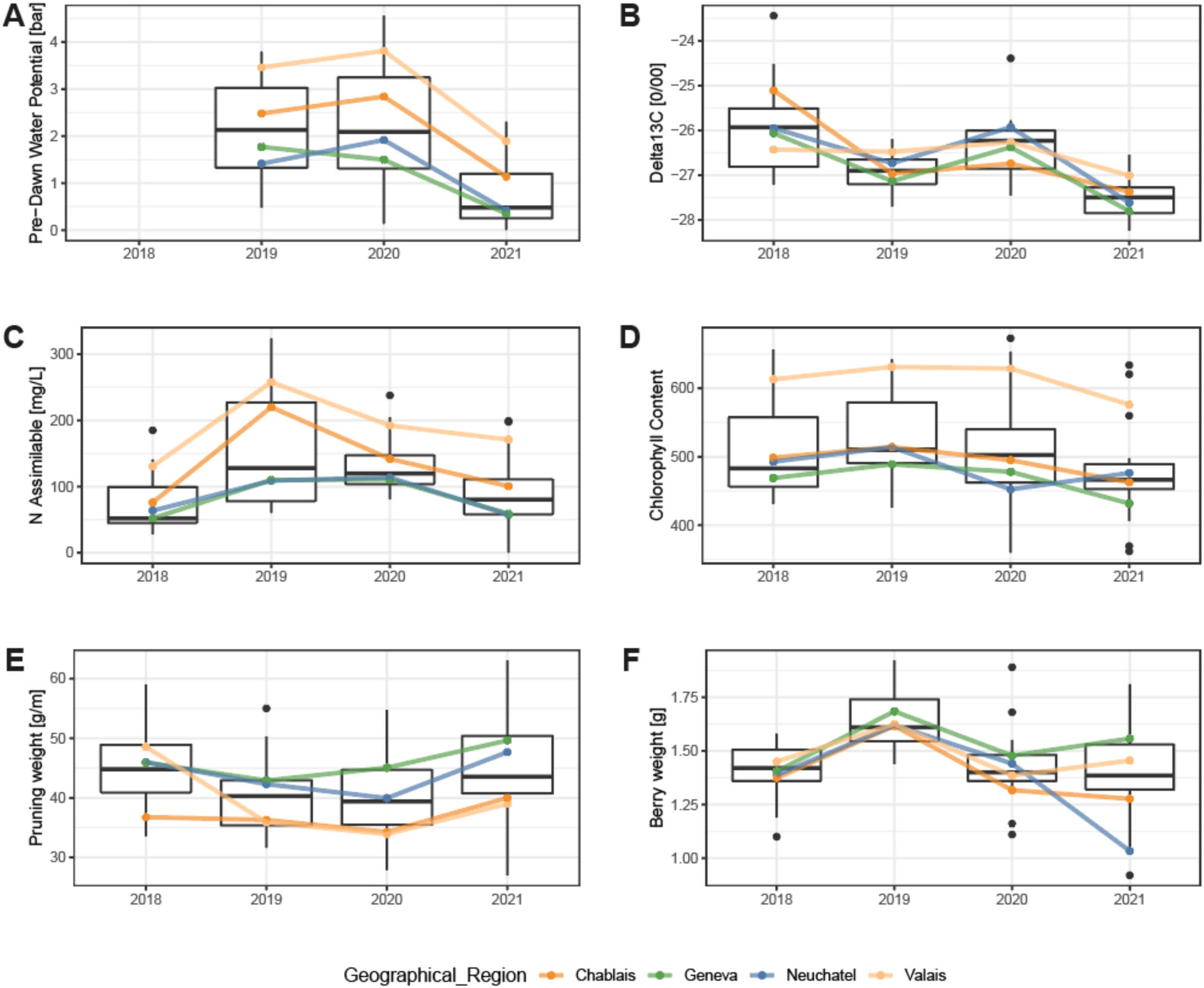
Mean values of measured physiological indexes of plots categorized by geographical region across four consecutive years (2018-2021) **A** pre-dawn leaf water potential (Ψ_PD_), **B** δ13C, **C** Assimilable nitrogen, **D** Chlorophyll content index (N-tester), **E** Pruning weight, **F** Berries weight.

