## Supplementary figures and images for "A systemic approach allows to identify the pedoclimatic conditions most critical in the susceptibility of a grapevine cultivar to esca/Botryosphaeria dieback"

### Supplementary Tables and Figures

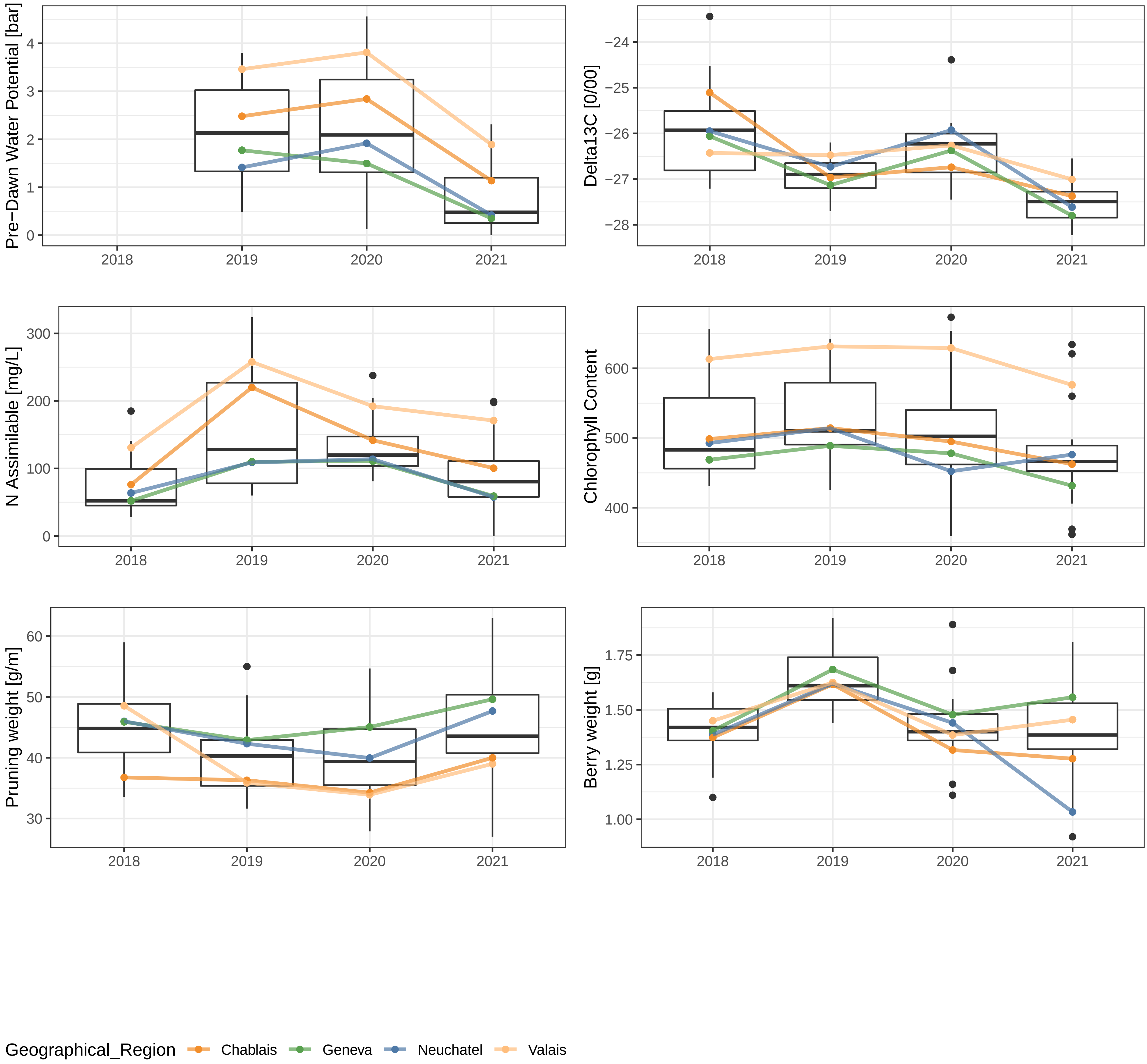
